# Human blood plasma proteins modeling and binding affinities with Δ^9^-tetrahydrocannabinol active metabolites: *In silico* approach

**DOI:** 10.1101/2022.04.13.488197

**Authors:** Shravan B. Rathod, Jinal C. Soni, Priyanshu Verma, Yogita Rawat, Neha Periwal, Pooja Arora, Vikas Sood, Mohmedyasin F. Mansuri

## Abstract

Tetrahydrocannabinol (THC) is a key psychotropic constituent of *cannabis sativa*. It is also known as Δ^9^-tetrahydrocannabinol (Δ^9^-THC). Previous study suggested that owing to its high lipophilicity, it piles up in adipose tissue and it is disseminated into blood stream for prolonged time. Research suggests that numerous diseases such as multiple sclerosis, neurodegenerative disorders, epilepsy, schizophrenia, osteoporosis, cancer, glaucoma and cardiovascular disorders can be treated using this substance. However, apart from having therapeutic potential, many studies have reported detrimental outcomes along with addiction of Δ^9^-THC for short-term and long-term consumption. Thus, in this study, we determined the binding affinities of Δ^9^-THC and its two active metabolites, 11-Hydroxy-Δ^9^-tetrahydrocannabinol (11-OH-Δ^9^-THC) and 8beta,11-dihydroxy-Δ^9^-tetrahydrocannabinol (8β,11-diOH-Δ^9^-THC) with 401 human blood plasma proteins using molecular docking analysis. Results show that Δ^9^-THC has greater binding potential with plasma proteins as compared to other two metabolites. Overall, ADGRE5, ALB, APOA5, APOD, CP, PON1 and PON3 proteins showed the highest binding affinities with three cannabis metabolites.

## Introduction

The consumption of cannabis in teenagers has surged significantly and, its adverse effects on mental health and political debate on whether make it legalise or not need urgent molecular level study of cannabis metabolites [1]. Even though epidemiological studies reported the risk of overconsumption of cannabis during adolescence leads to neuropsychiatric disorders in their later life, it still remains widely used drug illicit [2–4]. Researchers have shown that administration of a key ingredient, Δ^9^-tetrahydrocannabinol (Δ^9^-THC) of cannabis in animals during their adolescence, induced biochemical and behavioural signs of psychosis and depression later in life [5–10].

Interestingly, it was noted that the exposure of THC to female rats during adolescence is linked with epigenetic modification of histone H3 in Prefrontal cortex (PFC). Specifically, trimethylation at Lys9 position in N-terminus of histone H3. This alternation in histone H3 causes impacts on genes which are closely linked to the neuroplasticity and Endocannabinoid system (ECS) mechanisms that ultimately leads to diseases. However, THC exposure to adults results into minor effects on epigenome[1]. Thus, these findings suggest that the risks of neuropsychiatric disorders, CNS alternation and the cognitive impairment are escalated during adulthood due to the consumption of cannabis in adolescence [11]. Another study revealed that Δ^9^-THC activates the presynaptic Cannabinoid receptor type 1 (CB1) receptor which is key player in CNS development during prenatal, postnatal and adolescence periods [12–14]. The CB1 is G protein-coupled receptors (GPCR) and found in glial cells and neurons and cells [15]. Another, Cannabinoid receptors type 2 (CB2) is also GPCR which is generally located in hematopoietic cells and some specific regions of the brain and peripheral cells have CB2 receptor. CB2 plays an important role in immunity pathways and its activation leads to immunomodulatory and anti-inflammatory response. CB1 pathways are activated through the stimulation of CB2 [16,17]. Further, Abey N. O. investigated the effects of Δ^9^-THC on blood chemistry and organs cytoarchitecture and, it was observed significant decline in brain cognitive function, brain total proteins, and nitric oxide. Additionally, statistically significant difference were observed in various tissues and blood plasma [18]. Furthermore, computational and in vitro studies of Δ^9^-THC unveiled that this constituent strongly binds to three Fatty acid-binding proteins (FABPs), FABP3, FABP5 and FABP7 [19]. According to Schenk S. et al. analysis, they have validated and reported that there are 1193 different proteins are present in human blood plasma [20]. Literature review suggests that significant research has not been yet done on the binding profile of Δ^9^-THC and its metabolites with human blood plasma proteins. Hence, in this study, we determined the binding affinities of Δ^9^-THC and its two active metabolites, 11-OH-Δ^9^-THC and 8β,11-diOH-Δ^9^- THC with 401 plasma proteins using molecular docking approach. Fig. 1 shows the chemical structures of these three compounds.

**Fig. 1.**
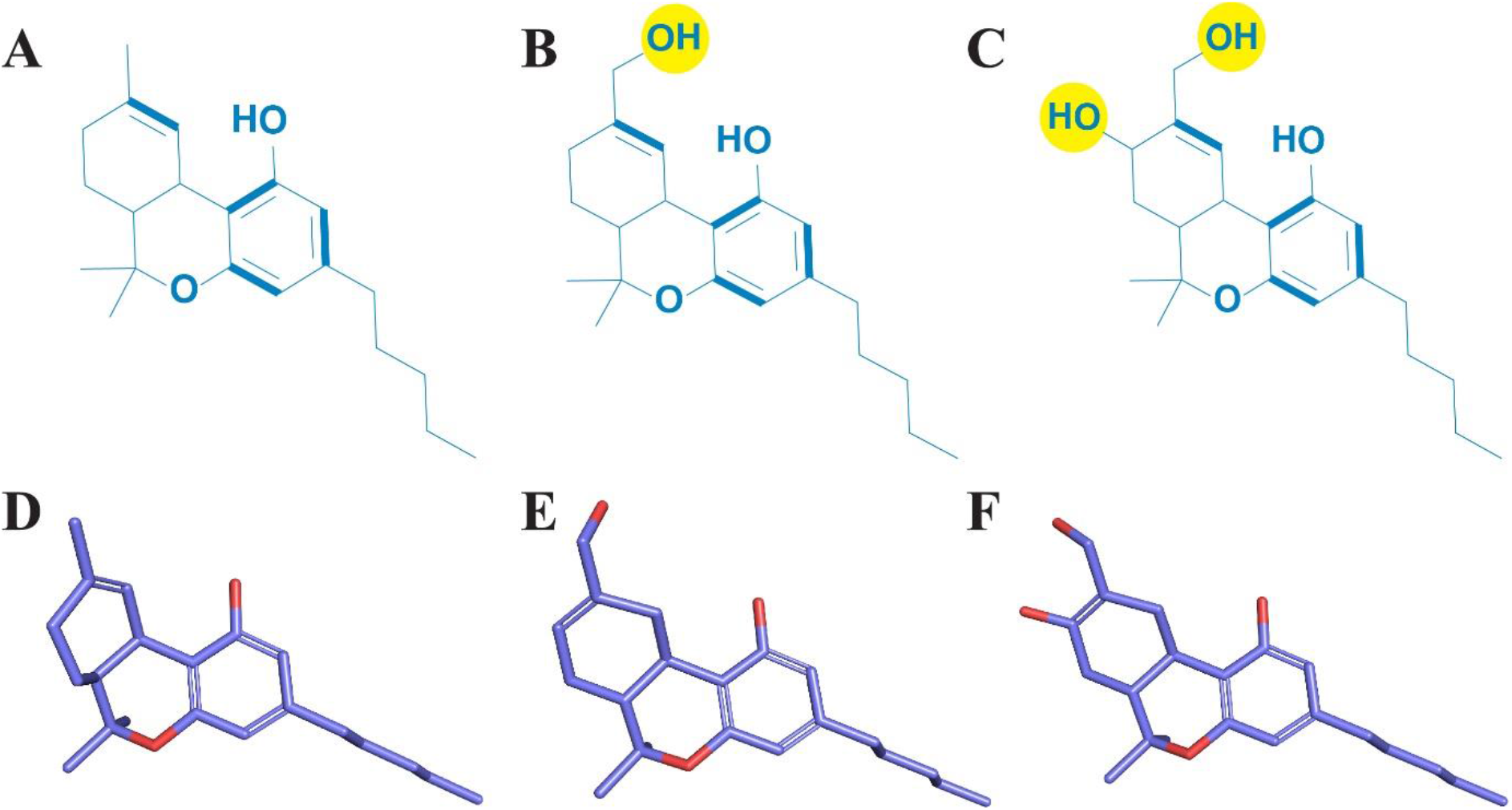
(A-C) 2D structures of Δ^9^-THC, 11-OH-Δ^9^-THC and 8β,11-diOH-Δ^9^-THC respectively. (D-F) 3D structures of Δ^9^-THC, 11-OH-Δ^9^-THC and 8β,11-diOH-Δ^9^-THC respectively. The yellow spheres indicate the addition of new hydroxyl groups to the Δ^9^-THC.

## Materials and methods

### Protein modeling and ligand preparation

Initially, the database of human blood plasma proteins detected by immunoassay was searched from The Human Protein Atlas (https://www.proteinatlas.org/humanproteome/blood+protein/proteins+detected+by+immunoassay) web portal. The total number of proteins were 419 on the database. Many protein structures were available on the Protein data bank (https://www.rcsb.org/) but large number of proteins had missing residues thus, we employed newly developed neural network-based RoseTTAFold [21] protein prediction tool (https://robetta.bakerlab.org/). This web tool has limitation that it accepts only proteins which have residues range between 26 and 1201. Thus, out of 419 plasma proteins, 18 proteins were excluded due to the lack of structure prediction. These 401 predicted structures were downloaded and further used to perform molecular docking analysis. The PDB structures of 401 human blood plasma proteins are given in Supplementary file 1.

In case of ligands, we considered Δ^9^-tetrahydrocannabinol (Δ^9^-THC: PubChem CID-16078) and its two active metabolites, 11-Hydroxy-Δ^9^-tetrahydrocannabinol (11-OH-Δ^9^-THC: PubChem CID-37482) and 8,11-dihydroxy-Δ^9^-tetrahydrocannabinol (8β,11-diOH-Δ^9^-THC: PubChem CID-126961369). Firstly, 3D structures of these ligands were retrieved from the PubChem Database (https://pubchem.ncbi.nlm.nih.gov/) in sdf format. Then, for the energy minimization, geometry optimization was performed on these ligands using Auto Optimization Tool available in Avogadro software [22]. The optimization was carried out by employing MMFF94s [23] force field and Steepest descent [24] algorithm. Finally, the structures were saved as mol2 format for further docking study.

### Molecular docking

To determine the binding affinities of three THC metabolites with 401 human blood plasma proteins, we utilized protein cavity-find guided Autodock Vina-based blind docker, CB-DOCK [25] web server available at http://clab.labshare.cn/cb-dock/php/. To calculate the binding affinity (Docking score) of these three metabolites with 401 plasma proteins, protein as a PDB and ligand as a mol2 format were uploaded to the server. We ran total 1203 (3*401) jobs at the server. Initially, this tool searches top five binding cavities inside the protein and orders them on the basis of cavity size. Then, it starts docking of ligand with receptor and calculate the binding affinity for best pose inside each cavity. Additionally, server provides details of cavity (volume, centre and size coordinates) along with binding affinity (Vina score). Further, the interactions analysis was carried out using Discovery Studio v.20.1 tool and UCSF Chimera v.1.15 was used to make figures of top two complexes. The Electrostatic potential (ESP) surface analysis of proteins was performed using PyMOL APBS Electrostatics plugin.

## Results and discussion

Molecular docking is a powerful computational approach to make drug discovery processes fast and less expensive. Molecular docking has many success stories for the designing inhibitors against HIV-1, cancer and bacterial targets [26–29]. Hence, for the primary investigation of binding affinities of Δ^9^-THC and its two active metabolites (11-OH-Δ^9^-THC and 8β,11-diOH-Δ^9^-THC) with human blood plasma proteins, we used blind docking approach.

Docking results show that binding affinities of Δ^9^-THC, 11-OH-Δ^9^-THC and 8β,11-diOH-Δ^9^-THC with 401 human blood plasma proteins vary between -5.0 and -10.5 kcal/mol. However, Δ^9^-THC showed the greater binding affinities as compared to other two metabolites (Fig. 2). This indicates that plasma proteins have overall hydrophobic nature as they have highest binding affinity with Δ^9^- THC. Δ^9^-THC is less polar (more hydrophobic) than 11-OH-Δ^9^-THC and 8β,11-diOH-Δ^9^-THC (Fig. 1). The binding affinities of all these three metabolites are given in Supplementary Table S1. Binding affinities of top 20 complexes are shown in Table 1.

**Table 1.** Binding affinities of top 20 complexes of Δ^9^-THC and its two metabolites (Ligands) with human blood plasma proteins.

| Sr.<br>No. | Ligands |  |  |  |  |  |
| --- | --- | --- | --- | --- | --- | --- |
| | $\Delta^9$-THC | | 11-OH-$\Delta^9$-THC | | 8β,11-diOH-$\Delta^9$-THC | |
|  | Protein | Binding<br>affinity<br>(kcal/mol) | Protein | Binding<br>affinity<br>(kcal/mol) | Protein | Binding<br>affinity<br>(kcal/mol) |
| 1 | APOA5 | -10.3 | ADGRE5 | -9.6 | APOA5 | -9.3 |
| 2 | APOD | -10.1 | CP | -9.5 | ADGRE5 | -9.1 |
| 3 | PON1 | -9.5 | ALB | -8.9 | APOD | -9.1 |
| 4 | CP | -9.3 | APOA5 | -8.9 | CP | -9.1 |
| 5 | ANGPTL3 | -9.2 | APOD | -8.9 | PON3 | -9.1 |
| 6 | LEPR | -9.1 | B2M | -8.9 | VTN | -8.9 |
| 7 | ADGRE5 | -9 | CETP | -8.9 | ALB | -8.8 |
| 8 | FGG | -8.9 | VTN | -8.9 | B2M | -8.8 |
| 9 | AFM | -8.8 | BPI | -8.8 | BPI | -8.7 |
| 10 | PLA2G2A | -8.8 | AFP | -8.6 | NAMPT | -8.7 |
| 11 | APOF | -8.7 | LBP | -8.6 | SELE | -8.6 |
| 12 | PDGFB | -8.7 | LEPR | -8.6 | FGG | -8.5 |
| 13 | PON3 | -8.7 | NAMPT | -8.6 | HPX | -8.5 |
| 14 | ALB | -8.6 | PLTP | -8.6 | PON1 | -8.5 |
| 15 | BMP6 | -8.6 | IL1RL1 | -8.5 | BMP6 | -8.4 |
| 16 | C9 | -8.6 | OSM | -8.5 | FABP3 | -8.4 |
| 17 | IL1RL1 | -8.6 | PON1 | -8.5 | HABP2 | -8.4 |
| 18 | SELE | -8.6 | BMP7 | -8.4 | LBP | -8.4 |
| 19 | VTN | -8.6 | FABP3 | -8.4 | MMP9 | -8.4 |
| 20 | BPI | -8.5 | FCRL5 | -8.4 | OSM | -8.4 |

**Fig. 2.**
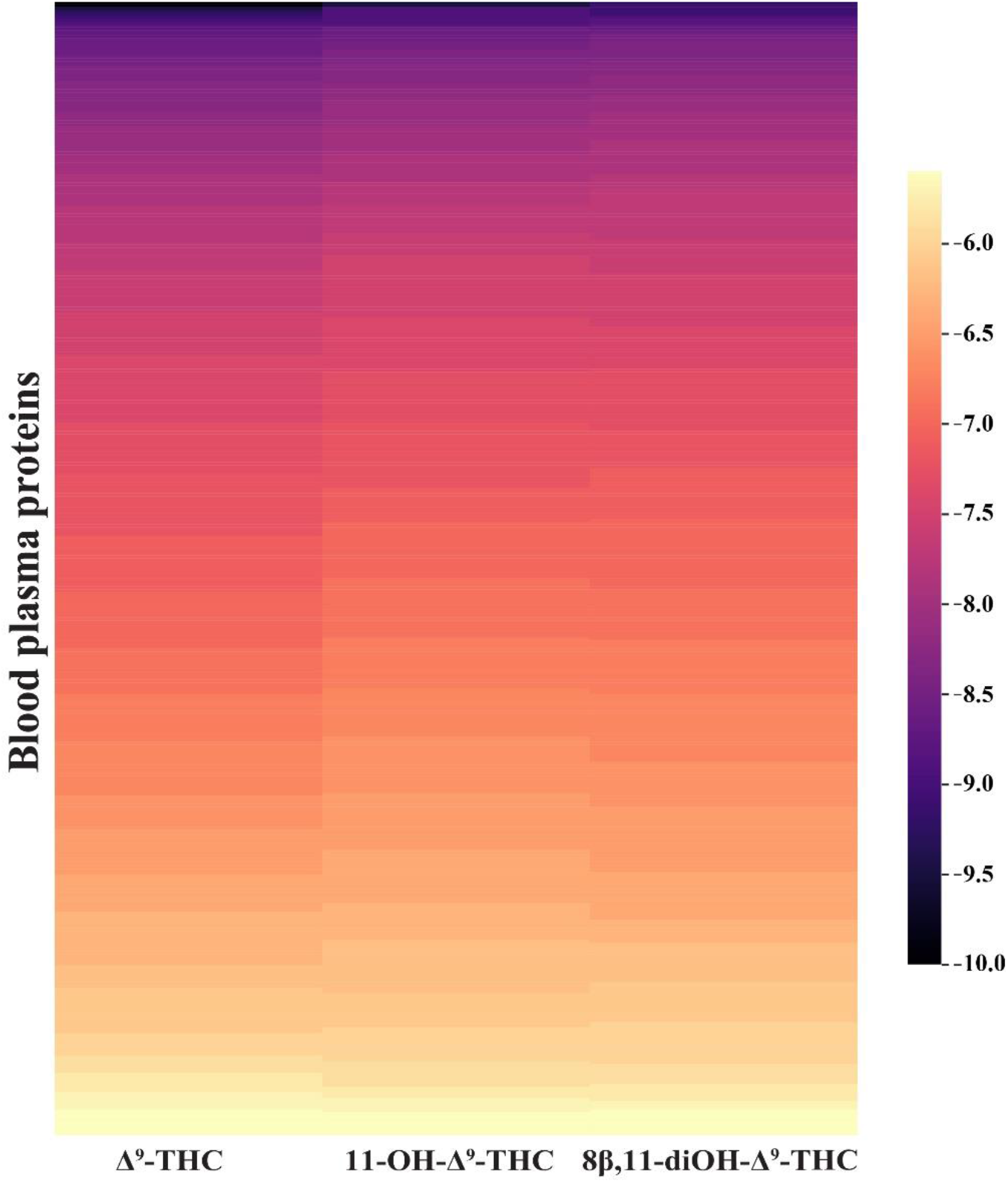
The heatmap of binding affinity (Vina score in kcal/mol) of Δ^9^-THC and its two metabolites with 401 human blood plasma proteins. Binding affinities correspond to color bar.

Additionally, we have carried out interaction analysis of top-two complexes of each metabolite. In case of Δ^9^-THC, Apolipoprotein A5 (APOA5) & Apolipoprotein D (APOD) have the large value of docking score, -10.3 & -10.1 kcal/mol respectively (Table 1). The structure and interaction plot of APOA5:Δ^9^-THC and APOD:Δ^9^-THC complexes are illustrated in Fig. 3. It has been reported that APOA5 plays vital role to metabolize triglyceride and triglyceride-rich lipoproteins. Current findings suggest the link between APOA5 and obesity [30]. Apolipoprotein D (APOD) is structurally different from the other apolipoproteins and it is lipocalin family member. It is widely expressed in glial cells and neurons of peripheral and central nervous system and carrier of tiny lipophilic candidates. It is not only transporter but has vital role in modulating the oxidation condition and stability of lipophilic molecules. Previous studies suggest that APOD was upregulated in stroke, schizophrenia and Alzheimer’s disease conditions. Thus, it has neuroprotective functions [31]. APOD forms barrel with eight antiparallel β-strands and has α-helices at the side. It has around ∼15 Å deep hydrophobic cavity and entry diameter is also 15 Å (Fig. 3C) [32].

**Fig. 3.**
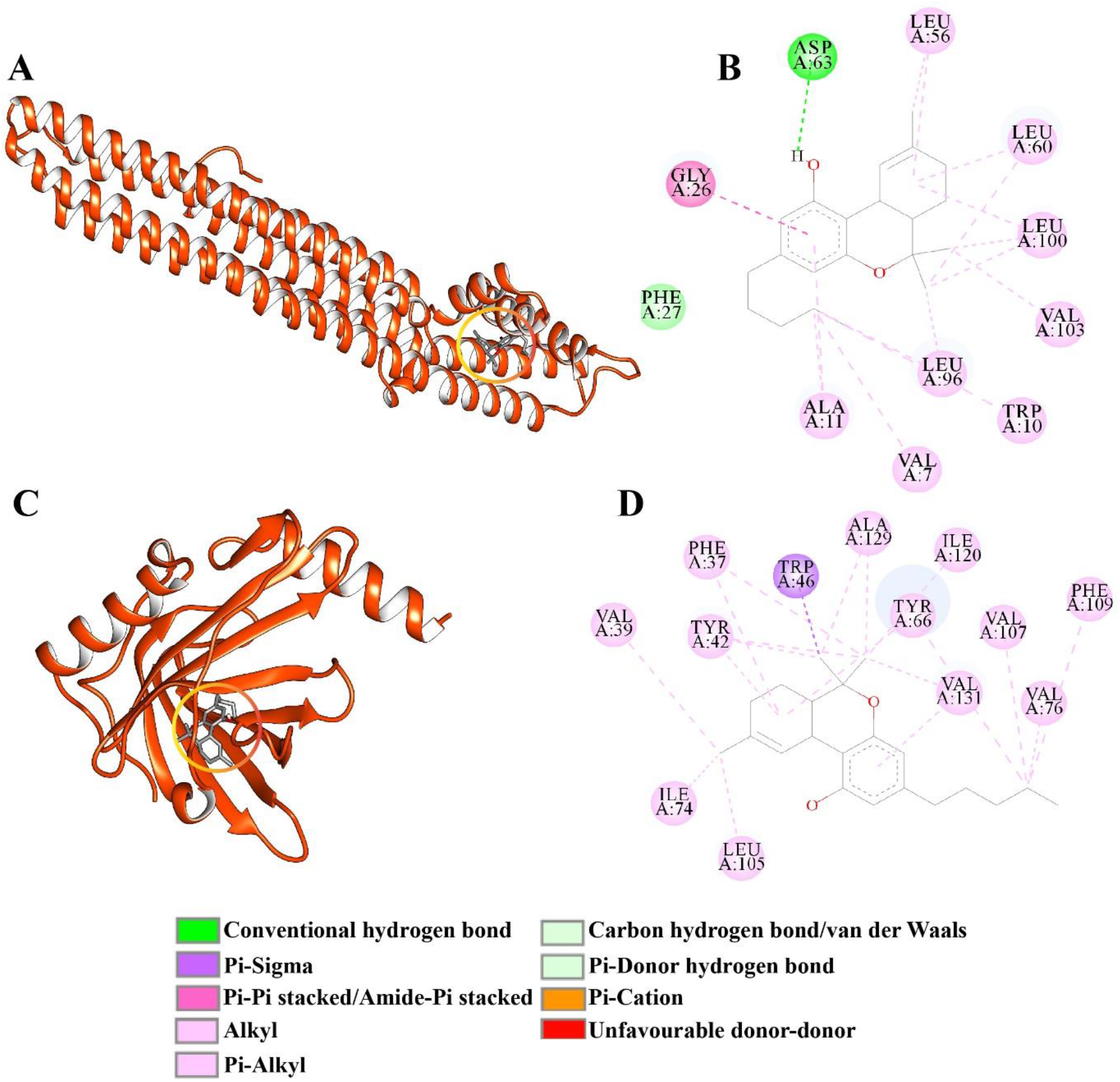
Complex structures and their 2D interactions. (A) APOA5:Δ^9^-THC complex. (B) APOA5-Δ^9^-THC interactions. (C) APOD:Δ^9^-THC complex. (D) APOD:Δ^9^-THC interactions. (APOA5: Apolipoprotein A5 & APOD: Apolipoprotein D). Orange-red circle shows the ligand position.

Fig. 3A & 3C represent the structure of APOA5:Δ^9^-THC and APOD:Δ^9^-THC complexes. It can be seen from the Fig. 3B that in APOA5:Δ^9^-THC complex, Val7, Trp10, Ala11, Gly26, Phe27, Leu56, Leu60, Leu96, Leu100 and Val103 residues have hydrophobic interactions such as π-alkyl and alkyl-alkyl. A single hydrogen bond is formed by Asp63 and hydroxyl group of Δ^9^-THC. Whereas, in APOD:Δ^9^-THC complex, Val, Phe, Leu, Tyr and Ile are the common residues which have hydrophobic interaction with ligand (Fig. 3D). The residues from the APOD interact with Δ^9^-THC are similar to the previously reported catalytic site residues. However, the number of residues interacting with Δ^9^-THC in APOD:Δ^9^-THC complex is slightly higher as compared to residues in APOA5:Δ^9^-THC complex.

Considering 11-OH-Δ^9^-THC metabolite, it can be observed (Table 1) that Adhesion G protein-coupled receptor E5 (ADGRE5) and Ceruloplasmin (CP) have better affinities around -9.5 kcal/mol in comparison with other proteins. The ADGRE5 is the subclass of G-protein coupled receptors (GPCRs) that has pharmacological significance [33]. It is also known as Cluster of differentiation 97 (CD97). This protein is broadly expressed muscle, immune, stem and progenitor cells [34-36]. Additionally, recent study reported that ADGRE5/CD97 helps to suppress Nuclear factor kappa B (NF-κB) through upregulating the Peroxisome proliferator-activated receptor gamma (PPAR-γ) in leukocytes [37]. Some studies also revealed that CD97 was found overexpressed in various tumours and attunes tumorigenesis [38].

The Ceruloplasmin (CP) belongs to oxidase family and copper enzyme present in serum [39]. In neurodegenerative diseases such as Parkinson’s and Alzheimer’s, CP was found in Cerebrospinal fluid (CSF) with modified structure and state [40]. Like APOD, CP has also neuroprotective role [41]. In CP, six cupredoxin domains are arranged side-by-side and form a cavity. CP has trinuclear copper cluster and mononuclear copper centre with a 12-13 Å distance between them. The trinuclear copper cluster is formed by three copper ions which are located in three even domains. This cluster has also one type II (T2) and two type III (T3) copper ions [42].

Complex structures of 11-OH-Δ^9^-THC with ADGRE5 and CP are given in Fig. 4A and C respectively. While the interactions maps are shown in Fig. 4B and D respectively. In ADGRE5 complex, none of the hydroxyl groups have polar interactions with surrounding residues. All interactions are hydrophobic in nature. Phe623 and Phe771 have π-π stacking interactions with aromatic ring in ligand. Remaining interactions come from hydrophobic amino acids such as Phe597, Leu686, Leu697, Val700, Leu757, Ile759 and Phe760 (Fig. 4B). However, 11-OH-Δ^9^-THC has polar interactions with CP. Lys288 forms hydrogen bond with hydroxyl group. Arg649 interacts with ligand through π-cation interaction. Phe659, Leu664 and Tyr986 have hydrophobic (π-alkyl & alkyl-alkyl) interactions (Fig. 4D). Additionally, Asn287 has unfavourable contact with hydroxyl group of aromatic ring due to its donor nature and steric repulsion.

**Fig. 4.**
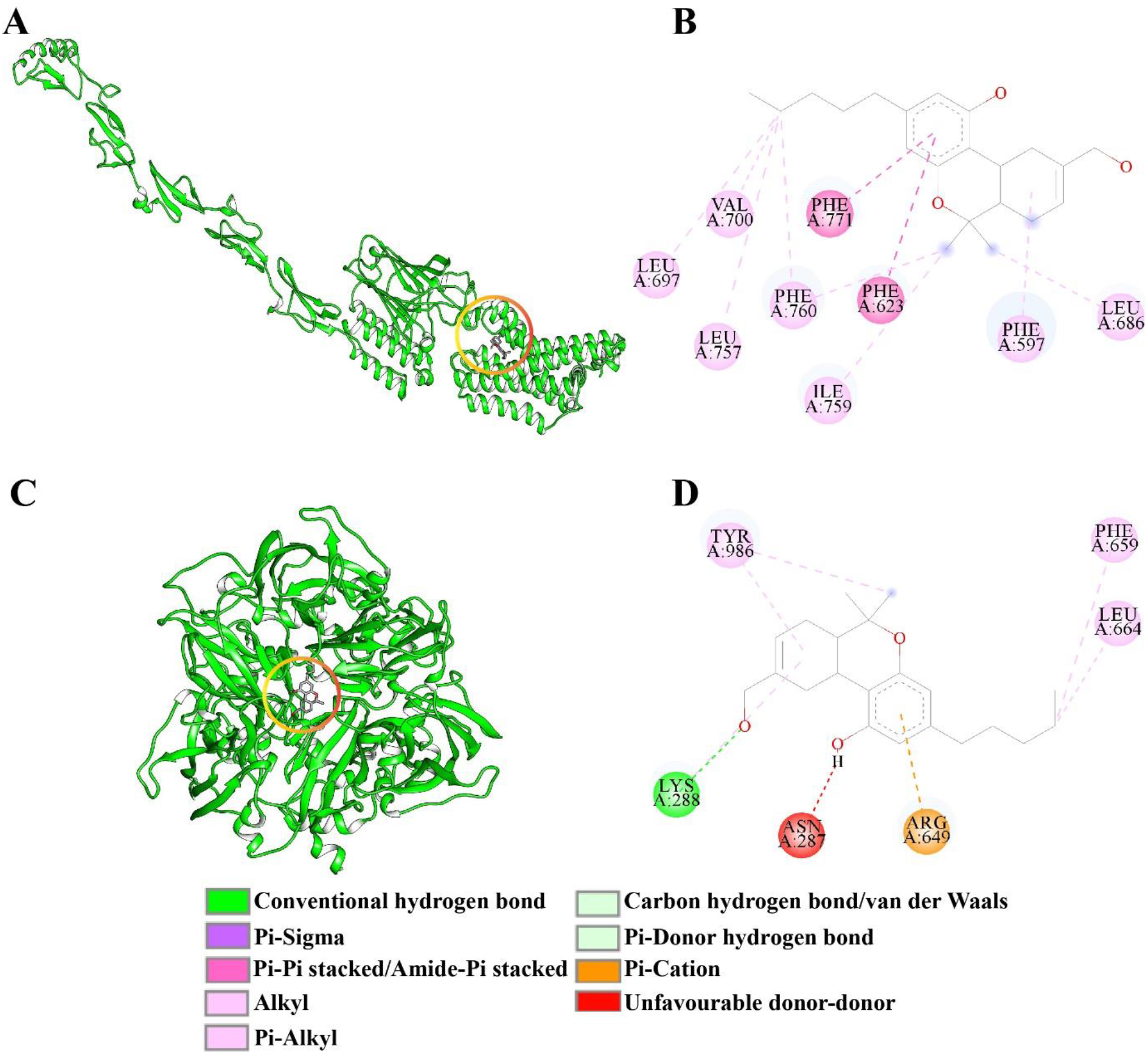
Complex structures and their 2D interactions. (A) ADGRE5:11-OH-Δ^9^-THC complex. (B) ADGRE5:11-OH-Δ^9^-THC interactions. (C) CP:11-OH-Δ^9^-THC complex. (D) CP:11-OH-Δ^9^-THC interactions. (ADGRE5: Adhesion G protein-coupled receptor E5 & CP: Ceruloplasmin). Orange-red circle shows the ligand position.

Finally, APOA5 and ADGRE5 were observed to have the large binding affinities (∼-9.0 kcal/mol) with third metabolite, 8β,11-diOH-Δ^9^-THC. The structures of APOA5:8β,11-diOH-Δ^9^-THC and ADGRE5:8β,11-diOH-Δ^9^-THC complexes are illustrated in Fig. 5A and 5C respectively whereas their interactions maps are shown in Fig. 5B and 5D respectively. In these complexes, hydrophobic interactions have been found dominant but both have polar interactions also. In APOA5:8β,11-diOH-Δ^9^-THC, Lys48 and Asp63 are forming hydrogen bonds with the hydroxyl groups of ligand. And, Trp10, Ala11, Leu14, Tyr30 and Leu96 interact through alkyl-alkyl hydrophobic interactions (Fig. 5B). Also, it has a single unfavourable interaction with Asp63. But, in ADGRE5:8β,11-diOH-Δ^9^-THC, the hydrogen bond is formed between Asn775 and aromatic hydroxyl group and, carbon-hydrogen (C-H) polar hydrogen bond between Thr772 and methylene group carbon where hydroxyl group is attached (Fig. 5D). Two aromatic residues, Phe623 and Phe771 show π-π stacking interactions with aromatic ring of ligand. And, large contribution comes from hydrophobic residues, Leu686, Leu697, Val700, Ile759 and Phe760 (Fig. 5D).

**Fig. 5.**
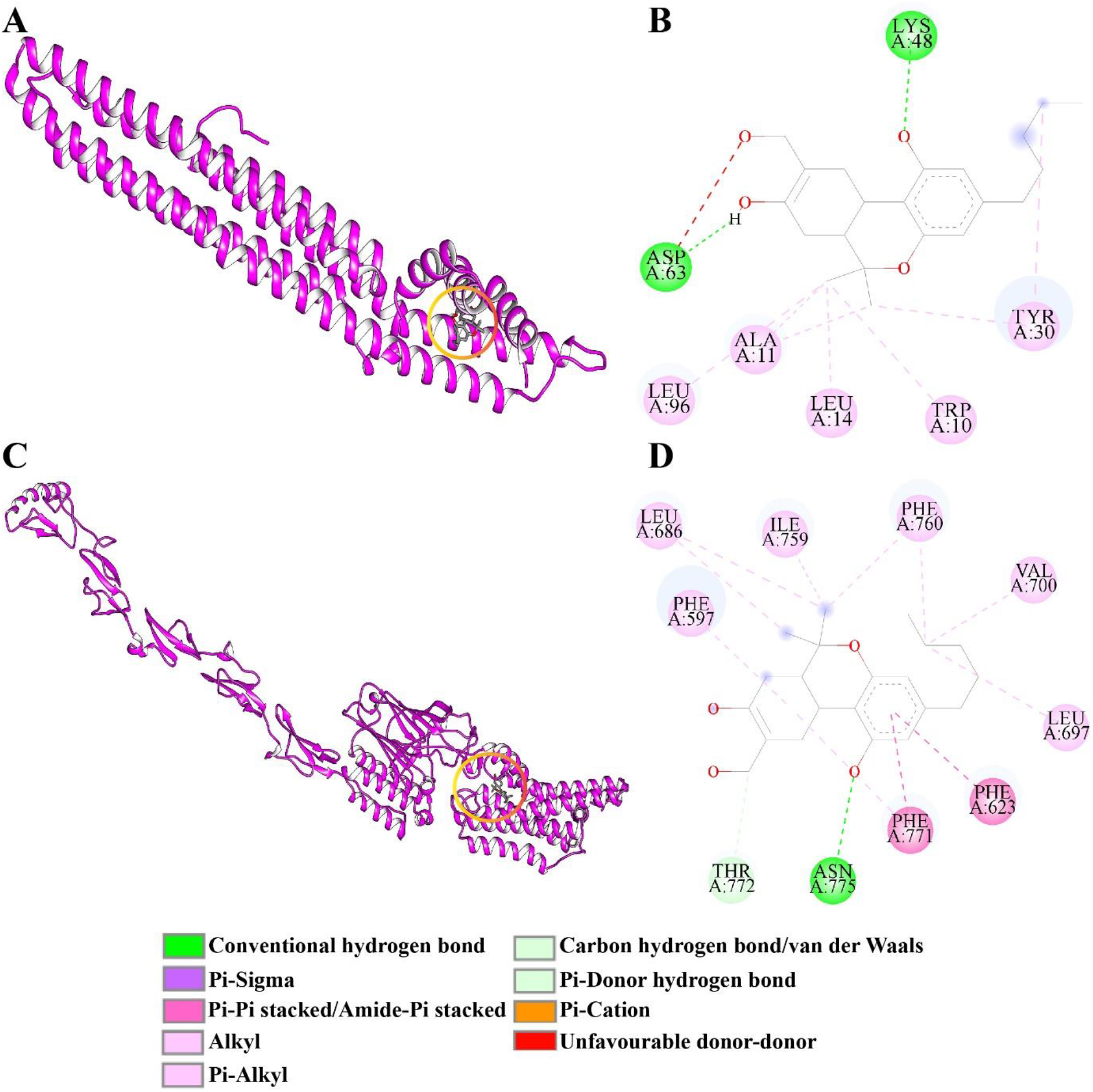
Complex structures and their 2D interactions. (A) APOA5:8,11-diOH-Δ^9^-THC complex. (B) APOA5-8,11-diOH-Δ^9^-THC interactions. (C) ADGRE5:8,11-diOH-Δ^9^-THC complex. (D) ADGRE5:8,11-diOH-Δ^9^-THC interactions. (APOA5: Apolipoprotein A5 & ADGRE5: Adhesion G protein-coupled receptor E5). Orange-red circle shows the ligand position.

Furthermore, we have also carried out ESP analysis for APOA5, APOD, ADGRE5 and CP to probe the ligand binding cavity electrostatic properties. Fig. 6A-D illustrate the hydrophobic nature of cavities in each four plasma proteins, APOA5, APOD, ADGRE5 and CP respectively.

**Fig. 6.**
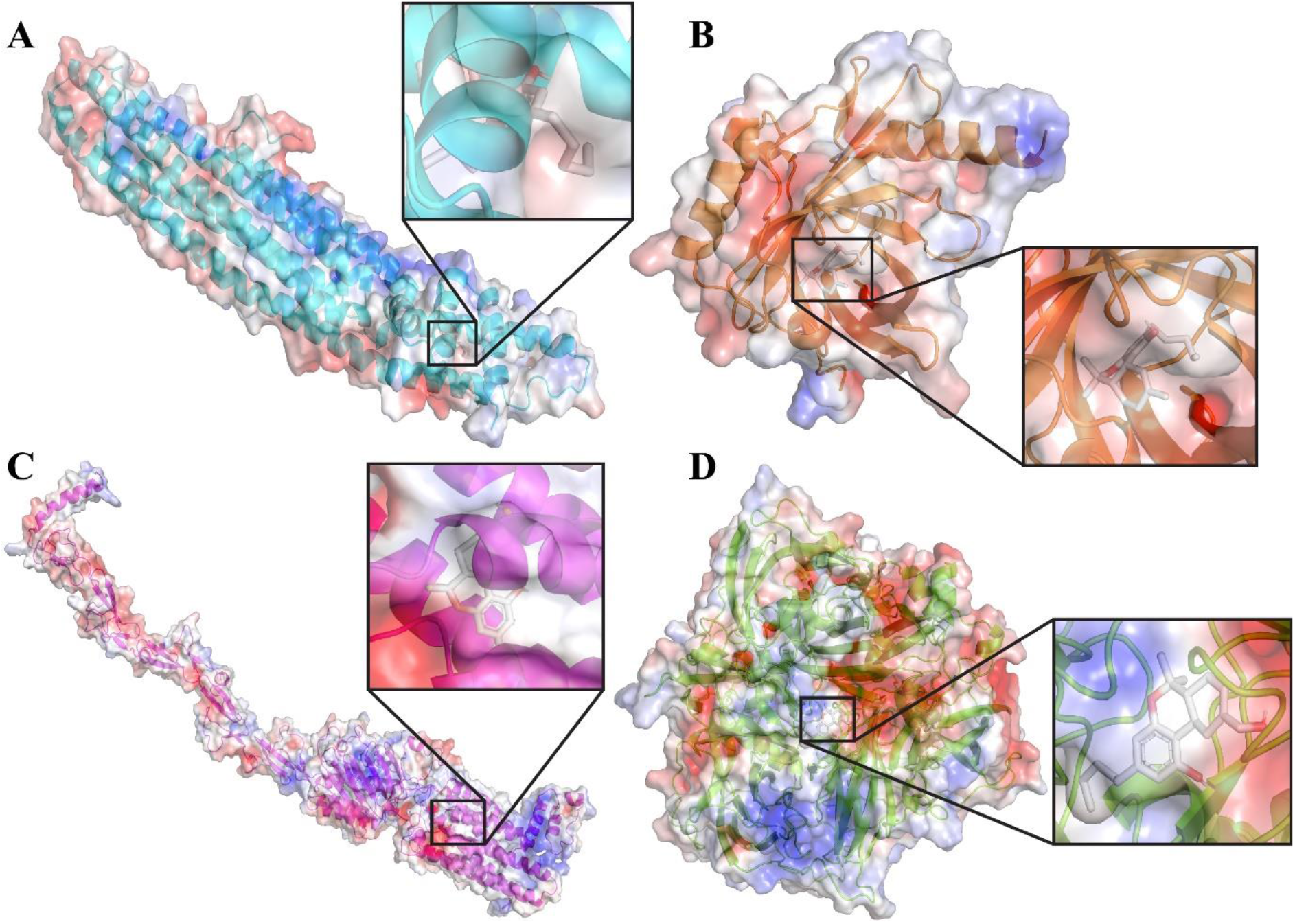
Electrostatic potential surface (ESP) of four proteins. (A) Apolipoprotein A5 (APOA5). (B) Apolipoprotein D (APOD). (C) Adhesion G protein-coupled receptor E5 (ADGRE5). (D) Ceruloplasmin (CP). Ligand position is indicated and zoomed in black square box.

## Conclusion

The principal constituent of cannabis, Δ^9^-THC has therapeutic significance as well as some obscurity in adverse impacts on health. Owing to the lipophilic nature of Δ^9^-THC, it can bind various human blood plasma proteins. Hence, we investigated the bindings of Δ^9^-THC and its two active metabolites (11-OH-Δ^9^-THC & 8β,11-diOH-Δ^9^-THC) with 401 human blood plasma proteins using blind docking CB-DOCK tool. Further, we performed interactions analysis of top-two complexes of each constituent with plasma proteins. Results suggest that Δ^9^-THC showed higher tendency towards plasma protein binding as compared to its two metabolites. The blood plasma proteins such as, ADGRE5, ALB, APOA5, APOD, CP, PON1 and PON3 have suitable binding cavities to adapt Δ^9^-THC and its active metabolites in comparison with remaining plasma proteins. Our study is a primary investigation of binding affinity between Δ^9^-THC and plasma proteins. However, our study provides input to further probe the structural, functional and dynamic impacts of these metabolites on blood plasma proteins leading to adverse impacts of these cannabinoids on human health by performing further studies.

## Acknowledgements

SBR is thankful to his Chemistry Department for providing computational facilities and infrastructure.

## Graphical abstract

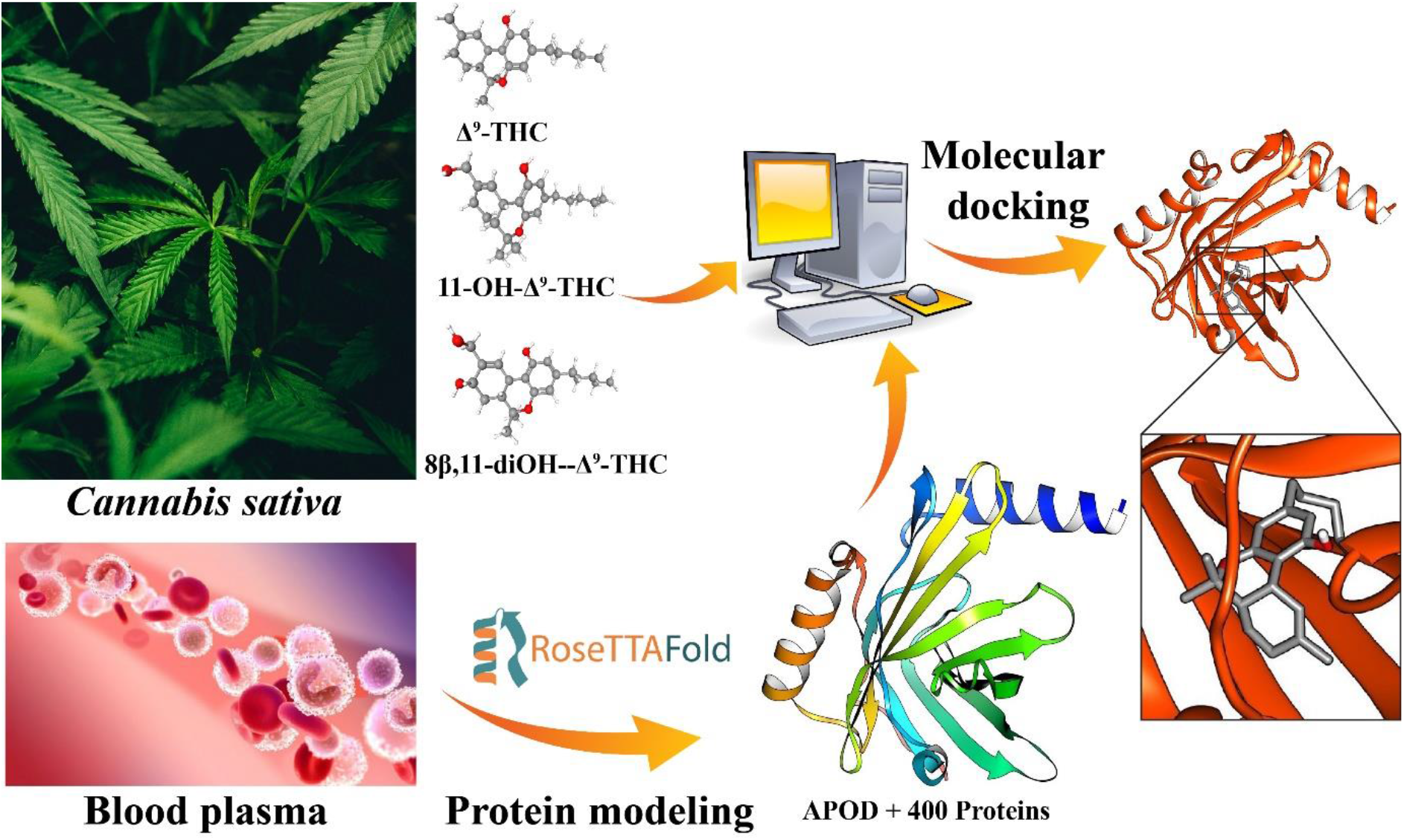

The sequences of the 401 human blood plasma proteins were retrieved from the UniProt database and RoseTTAFold protein structure prediction tool was employed to predict the structures of proteins. Next, three metabolites, Δ^9^-THC, 11-OH-Δ^9^-THC and 8β,11-diOH-Δ^9^-THC were downloaded from the PubChem database and energy minimization of these three metabolites were done using Avogadro molecular graphic tool. Finally, the molecular docking of these three substances was performed with 401 (3*401= 1203) using blind docker CB-DOCK tool and, interactions analysis of top two complexes of each metabolite was carried out.

## References

[1] P. Prini, F. Rusconi, E. Zamberletti, M. Gabaglio, F. Penna, M. Fasano, E. Battaglioli, D. Parolaro, T. Rubino, Adolescent THC exposure in female rats leads to cognitive deficits through a mechanism involving chromatin modifications in the prefrontal cortex, J. Psychiatry Neurosci. 43 (2018) 87–101. https://doi.org/10.1503/jpn.170082.

[2] S. Lev-Ran, M. Roerecke, B. Le Foll, T.P. George, K. McKenzie, J. Rehm, The association between cannabis use and depression: A systematic review and meta-analysis of longitudinal studies, Psychol. Med. 44 (2014) 797–810. https://doi.org/10.1017/S0033291713001438.

[3] M. Di Forti, H. Sallis, F. Allegri, A. Trotta, L. Ferraro, S.A. Stilo, A. Marconi, C. La Cascia, T.R. Marques, C. Pariante, P. Dazzan, V. Mondelli, A. Paparelli, A. Kolliakou, D. Prata, F. Gaughran, A.S. David, C. Morgan, D. Stahl, M. Khondoker, J.H. MacCabe, R.M. Murray, Daily use, especially of high-potency cannabis, drives the earlier onset of psychosis in cannabis users, Schizophr. Bull. 40 (2014) 1509–1517. https://doi.org/10.1093/schbul/sbt181.

[4] S.T. Wilkinson, R. Radhakrishnan, D.C. D’Souza, Impact of Cannabis Use on the Development of Psychotic Disorders, Curr. Addict. Reports. 1 (2014) 115–128. https://doi.org/10.1007/s40429-014-0018-7.

[5] T. Rubino, D. Vigano’, N. Realini, C. Guidali, D. Braida, V. Capurro, C. Castiglioni, F. Cherubino, P. Romualdi, S. Candeletti, M. Sala, D. Parolaro, Chronic Δ9-tetrahydrocannabinol during adolescence provokes sex-dependent changes in the emotional profile in adult rats: Behavioral and biochemical correlates, Neuropsychopharmacology. 33 (2008) 2760–2771. https://doi.org/10.1038/sj.npp.1301664.

[6] T. Rubino, N. Realini, D. Braida, T. Alberio, V. Capurro, D. Viganò, C. Guidali, M. Sala, M. Fasano, D. Parolaro, The Depressive Phenotype Induced in Adult Female Rats by Adolescent Exposure to THC is Associated with Cognitive Impairment and Altered Neuroplasticity in the Prefrontal Cortex, Neurotox. Res. 15 (2009) 291–302. https://doi.org/10.1007/s12640-009-9031-3.

[7] T. Rubino, P. Prini, F. Piscitelli, E. Zamberletti, M. Trusel, M. Melis, C. Sagheddu, A. Ligresti, R. Tonini, V. Di Marzo, D. Parolaro, Adolescent exposure to THC in female rats disrupts developmental changes in the prefrontal cortex, Neurobiol. Dis. 73 (2015) 60–69. https://doi.org/10.1016/j.nbd.2014.09.015.

[8] N. Realini, D. Vigano’, C. Guidali, E. Zamberletti, T. Rubino, D. Parolaro, Chronic URB597 treatment at adulthood reverted most depressive-like symptoms induced by adolescent exposure to THC in female rats, Neuropharmacology. 60 (2011) 235–243. https://doi.org/10.1016/j.neuropharm.2010.09.003.

[9] E. Zamberletti, S. Beggiato, L. Steardo, P. Prini, T. Antonelli, L. Ferraro, T. Rubino, D. Parolaro, Neurobiology of Disease Alterations of prefrontal cortex GABAergic transmission in the complex psychotic-like phenotype induced by adolescent delta-9-tetrahydrocannabinol exposure in rats, Neurobiol. Dis. 63 (2014) 35–47. https://doi.org/10.1016/j.nbd.2013.10.028.

[10] E. Zamberletti, M. Gabaglio, P. Prini, T. Rubino, D. Parolaro, Cortical neuroin flammation contributes to long-term cognitive dysfunctions following adolescent delta-9-tetrahydrocannabinol treatment in female rats, Eur. Neuropsychopharmacol. 25 (2015) 2404–2415. https://doi.org/10.1016/j.euroneuro.2015.09.021.

[11] J. Renard, M.O. Krebs, G. Le Pen, T.M. Jay, Long-term consequences of adolescent cannabinoid exposure in adult psychopathology, Front. Neurosci. 8 (2014) 1–14. https://doi.org/10.3389/fnins.2014.00361.

[12] V. Di Marzo, The endocannabinoid system: Its general strategy of action, tools for its pharmacological manipulation and potential therapeutic exploitation, Pharmacol. Res. 60 (2009) 77–84. https://doi.org/10.1016/j.phrs.2009.02.010.

[13] T. Harkany, M. Guzmán, I. Galve-Roperh, P. Berghuis, L.A. Devi, K. Mackie, The emerging functions of endocannabinoid signaling during CNS development, Trends Pharmacol. Sci. 28 (2007) 83–92. https://doi.org/10.1016/j.tips.2006.12.004.

[14] M. Ellgren, A. Artmann, O. Tkalych, A. Gupta, H.S. Hansen, S.H. Hansen, L.A. Devi, Y.L. Hurd, Dynamic changes of the endogenous cannabinoid and opioid mesocorticolimbic systems during adolescence: THC effects, Eur. Neuropsychopharmacol. 18 (2008) 826–834. https://doi.org/10.1016/j.euroneuro.2008.06.009.

[15] L.E. Klumpers, D.L. Thacker, A brief background on cannabis: From plant to medical indications, J. AOAC Int. 102 (2019) 412–420. https://doi.org/10.5740/jaoacint.18-0208.

[16] R.G. Pertwee, Receptors and channels targeted by synthetic cannabinoid receptor agonists and antagonists. Curr Med Chem. 17 (2010) 1360–1381. http://dx.doi.org/10.2174/092986710790980050

[17] E.D. Gonçalves, R.C. Dutra, Cannabinoid receptors as therapeutic targets for autoimmune diseases: where do we stand?, Drug Discov. Today. 24 (2019) 1845–1853. https://doi.org/10.1016/j.drudis.2019.05.023.

[18] N.O. Abey, Cannabis sativa (Marijuana) alters blood chemistry and the cytoarchitecture of some organs in Sprague Dawley rat models, Food Chem. Toxicol. 116 (2018) 292–297. https://doi.org/10.1016/j.fct.2018.04.023.

[19] M.W. Elmes, M. Kaczocha, W.T. Berger, K.N. Leung, B.P. Ralph, L. Wang, J.M. Sweeney, J.T. Miyauchi, S.E. Tsirka, I. Ojima, D.G. Deutsch, Fatty acid-binding proteins (FABPs) are intracellular carriers for Δ9-tetrahydrocannabinol (THC) and cannabidiol (CBD), J. Biol. Chem. 290 (2015) 8711–8721. https://doi.org/10.1074/jbc.M114.618447.

[20] S. Schenk, G.J. Schoenhals, G. de Souza, M. Mann, A high confidence, manually validated human blood plasma protein reference set, BMC Med. Genomics. 1 (2008). https://doi.org/10.1186/1755-8794-1-41.

[21] M. Baek, F. DiMaio, I. Anishchenko, J. Dauparas, S. Ovchinnikov, G.R. Lee, J. Wang, Q. Cong, L.N. Kinch, R. Dustin Schaeffer, C. Millán, H. Park, C. Adams, C.R. Glassman, A. DeGiovanni, J.H. Pereira, A. V. Rodrigues, A.A. Van Dijk, A.C. Ebrecht, D.J. Opperman, T. Sagmeister, C. Buhlheller, T. Pavkov-Keller, M.K. Rathinaswamy, U. Dalwadi, C.K. Yip, J.E. Burke, K. Christopher Garcia, N. V. Grishin, P.D. Adams, R.J. Read, D. Baker, Accurate prediction of protein structures and interactions using a three-track neural network, Science. 373 (2021) 871–876. https://doi.org/10.1126/science.abj8754.

[22] M.D. Hanwell, D.E. Curtis, D.C. Lonie, T. Vandermeersch, E. Zurek, G.R. Hutchison, Avogadro: an advanced semantic chemical editor, visualization, and analysis platform, J Cheminform. 4 (2012) 1–17. https://doi.org/10.1186/1758-2946-4-17.

[23] T.A. Halgren, Performance of MMFF94*, J. Comput. Chem. 17 (1996) 490–519. http://journals.wiley.com/jcc.

[24] C. Chen, Y. Huang, X. Ji, Y. Xiao, Efficiently finding the minimum free energy path from steepest descent path, J. Chem. Phys. 138 (2013) 1–9. https://doi.org/10.1063/1.4799236.

[25] Y. Liu, M. Grimm, W. tao Dai, M. chun Hou, Z.X. Xiao, Y. Cao, CB-Dock: a web server for cavity detection-guided protein–ligand blind docking, Acta Pharmacol. Sin. 41 (2020) 138–144. https://doi.org/10.1038/s41401-019-0228-6.

[26] R. Dayam, N. Neamati, Active site binding modes of the β-diketoacids: A multi-active site approach in HIV-1 integrase inhibitor design, Bioorganic Med. Chem. 12 (2004) 6371–6381. https://doi.org/10.1016/j.bmc.2004.09.035.

[27] A. Kazi, H. Lawrence, W.C. Guida, M.L. McLaughlin, G.M. Springett, N. Berndt, R.M.L. Yip, S.M. Sebti, Discovery of a novel proteasome inhibitor selective for cancer cells over non-transformed cells, Cell Cycle. 8 (2009) 1940–1951. https://doi.org/10.4161/cc.8.12.8798.

[28] N.A. Kahlous, M.A.M. Bawarish, M.A. Sarhan, M. Küpper, A. Hasaba, M. Rajab, Using Chemoinformatics, Bioinformatics, and Bioassay to Predict and Explain the Antibacterial Activity of Nonantibiotic Food and Drug Administration Drugs, Assay Drug Dev. Technol. 15 (2017) 89–105. https://doi.org/10.1089/adt.2016.771.

[29] C.R. Oliva, W. Zhang, C. Langford, M.J. Suto, C.E. Griguer, Repositioning chlorpromazine for treating chemoresistant glioma through the inhibition of cytochrome c oxidase bearing the COX4-1 regulatory subunit, Oncotarget. 8 (2017) 37568–37583. https://doi.org/10.18632/oncotarget.17247.

[30] X. Su, Y. Kong, D.Q. Peng, New insights into apolipoprotein A5 in controlling lipoprotein metabolism in obesity and the metabolic syndrome patients, Lipids Health Dis. 17 (2018) 1–10. https://doi.org/10.1186/s12944-018-0833-2.

[31] S. Dassati, A. Waldner, R. Schweigreiter, Apolipoprotein D takes center stage in the stress response of the aging and degenerative brain, Neurobiol. Aging. 35 (2014) 1632–1642. https://doi.org/10.1016/j.neurobiolaging.2014.01.148.

[32] A. Eichinger, A. Nasreen, J.K. Hyun, A. Skerra, Structural insight into the dual ligand specificity and mode of high density lipoprotein association of apolipoprotein D, J. Biol. Chem. 282 (2007) 31068–31075. https://doi.org/10.1074/jbc.M703552200.

[33] J. Hamann, G. Aust, D. Araç, F.B. Engel, C. Formstone, R. Fredriksson, R.A. Hall, B.L. Harty, C. Kirchhoff, B. Knapp, A. Krishnan, I. Liebscher, H.H. Lin, D.C. Martinelli, K.R. Monk, M.C. Peeters, X. Piao, S. Prömel, T. Schöneberg, T.W. Schwartz, K. Singer, M. Stacey, Y.A. Ushkaryov, M. Vallon, U. Wolfrum, M.W. Wright, L. Xu, T. Langenhan, H.B. Schiöth, International union of basic and clinical pharmacology. XCIV. adhesion G protein-coupled receptors, Pharmacol. Rev. 67 (2015) 338–367. https://doi.org/10.1124/pr.114.009647.

[34] M. Van Pel, H. Hagoort, J. Hamann, W.E. Fibbe, CD97 is differentially expressed on murine hematopoietic stem-and progenitor-cells, Haematologica. 93 (2008) 1137–1144. https://doi.org/10.3324/haematol.12838.

[35] G. Aust, E. Wandel, C. Boltze, D. Sittig, A. Schütz, L.C. Horn, M. Wobus, Diversity of CD97 in smooth muscle cells, Cell Tissue Res. 324 (2006) 139–147. https://doi.org/10.1007/s00441-005-0103-2.

[36] L.H. Jaspars, W. Vos, G. Aust, R.A.W. Van Lier, J. Hamann, Tissue distribution of the human CD97 EGF-TM7 receptor, Tissue Antigens. 57 (2001) 325–331. https://doi.org/10.1034/j.1399-0039.2001.057004325.x.

[37] S. Wang, Z. Sun, W. Zhao, Z. Wang, M. Wu, Y. Pan, H. Yan, J. Zhu, CD97/ADGRE5 Inhibits LPS Induced NF-B Activation through PPAR-γ Upregulation in Macrophages, Mediators Inflamm. 2016 (2016) 1–10. https://doi.org/10.1155/2016/1605948.

[38] W.Y. Tjong, H.H. Lin, The role of the RGD motif in CD97/ADGRE5-and EMR2/ADGRE2-modulated tumor angiogenesis, Biochem. Biophys. Res. Commun. 520 (2019) 243–249. https://doi.org/10.1016/j.bbrc.2019.09.113.

[39] N.E. Hellman, J.D. Gitlin, Ceruloplasmin metabolism and function, Annu. Rev. Nutr. 22 (2002) 439–458. https://doi.org/10.1146/annurev.nutr.22.012502.114457.

[40] A. Zanardi, M. Alessio, Ceruloplasmin deamidation in neurodegeneration: From loss to gain of function, Int. J. Mol. Sci. 22 (2021) 1–13. https://doi.org/10.3390/ijms22020663.

[41] B. Wang, X.-P. Wang, Does Ceruloplasmin Defend Against Neurodegenerative Diseases?, Curr. Neuropharmacol. 17 (2018) 539–549. https://doi.org/10.2174/1570159x16666180508113025.

[42] I. Bento, C. Peixoto, V.N. Zaitsev, P.F. Lindley, Ceruloplasmin revisited: Structural and functional roles of various metal cation-binding sites, Acta Crystallogr. Sect. D Biol. Crystallogr. 63 (2007) 240–248. https://doi.org/10.1107/S090744490604947X.

